# Minhee Analysis Package: An Integrated Software Package for Detection and Management of Spontaneous Synaptic Events

**DOI:** 10.1101/2021.05.27.443730

**Authors:** Yong Gyu Kim, Jae Jin Shin, Sang Jeong Kim

**Author notes:** Address for correspondence: Sang Jeong Kim, M.D., Ph.D. Department of Physiology, Seoul National University College of Medicine, 103 Daehangno, Jongro-gu, Seoul 03080, Republic of Korea.

## Abstract

To understand the information encoded in a connection between the neurons, postsynaptic current (PSC) has been widely measured as a primary index of synaptic strength in the field of neurophysiology. Although several automatic detection methods for PSCs have been proposed to simplify a workflow in the analysis, repetitive steps such as quantification and management of PSC data should be still performed with much effort. Here, we present Minhee Analysis Package, an integrated standalone software package that is capable of detecting, sorting, and quantifying PSC data. First, we developed a stepwise exploratory algorithm to detect PSC and validated our detection algorithm using the simulated and experimental data. We also described all the features and examples of the package so that users can use and follow them properly. In conclusion, our software package is expected to improve the convenience and efficiency of neurophysiologists to analyze PSC data by simplifying the workflow from detection to quantification. Minhee Analysis Package is freely available to download from http://www.github.com/parkgilbong/Minhee_Analysis_Pack.

## 1. INTRODUCTION

The neurons communicate with each other in a precisely timed manner within a complex neural network. To transfer information in real-time dialogues between them, they release and receive neurotransmitters via synapses, which is represented as synaptic transmission. In order to decode information in a neuronal synaptic connection, it is essential to detect, measure, and analyze the characteristics of synaptic transmission. Of several properties that characterize synaptic transmission, the postsynaptic current (PSC) has been extensively utilized to understand neuronal communication via synapses [1–3]. In terms of neurophysiology, PSCs are generated spontaneously or as a result of presynaptic spikes, which are represented as miniature or spontaneous PSC respectively. Therefore, quantitative analysis of PSCs in neurons is a fundamental step to characterize synaptic properties in the brain. While the results of quantitative analysis of PSCs are informative, it is time-consuming and laborious. To simplify this process in an efficient way, several automated methods have been suggested in the detection of spontaneous PSCs [4–10]. However, even after the reliable detection of PSCs, additional following steps still remain redundant for researchers to keep doing consecutive and iterative analysis to interpret the results. These steps include averaging and visualizing the data sorted by experimental conditions and then performing statistical assessments that are routinely used. Therefore, an integrated solution that includes quantification and management of PSC data would improve researcher’s method of analysis in a convenient and efficient way by simplifying the workflow from detection to quantification of PSC data.

Here, we present Minhee Analysis Package, an integrated software package that can detect and manage spontaneous synaptic events practically and precisely. It is built not just to detect spontaneous PSCs, but also to sort and visualize PSC data, and then perform the hypothesis testing, in a row. This package utilizes a stepwise exploratory algorithm to detect PSCs. In addition to the function of an event detection, users can briefly recall their result in the user's defined manner. With the successful retrieval of the data, the package automatically produces general and cumulative histograms to represent the distribution of data. The package also serves two hypothesis testing methods, an independent student’s t-test for comparing averaged values between the groups and Kolmogorov-Smirnov (K-S) test for comparing distribution of data between two groups. Variety of functions of Minhee Analysis Package are accessible via its easy-to-use graphical user interface (GUI). It can be downloaded at http://www.github.com/parkgilbong/Minhee_Analysis_Pack.

## 2. MATERIALS AND METHODS

### 2.1 Package development environment

The Minhee Analysis Package was developed by the LabVIEW (version 2017, National instrument, USA, RRID:SCR_014325), which includes a large library of functions for signal process and analysis. The LabVIEW development environment provides the application builder that enables the package to be distributed as a stand-alone application with a binary installer.

The Minhee Analysis Package took an advantage of built-in functions called VI (Virtual Instrument) in the LabVIEW development environment. The Savitzky-Golay (S-G) Filter VI^1^ is used for smoothing the recorded trace. The S-G filter performs a local polynomial regression around each point, and creates a smoothed value for each data point. It has two parameters to determine the degree of smoothness: the polynomial order and side points. We adopted these two parameters as adjustable parameters for PSCs event detection. The polynomial order determines the order of the polynomial for the local regression. The side points specify the number of data points in the moving window for local polynomial regression.

We also employ a built-in function of the LabVIEW advanced signal processing toolkit^2^ called Multiscale Peak Detection VI^3^ for the initial search step in the event detection algorithms. This function is utilized to detect peaks or valleys in a signal that are considered as local peaks or valleys in the initial search step of event detection. The value of the threshold parameter is set to 3, therefore, this function detects peaks or valleys above 3 pA in the signal.

To search for files that match users’ search pattern in the Minhee retriever, we utilize the Recursive File List VI^4^ function. Two features in the Minhee retriever, ***Search pattern*** and ***Folders to exclude***, have been directly adopted from this VI.

### 2.2 Simulation of postsynaptic currents (PSCs)

Simulated EPSCs were randomly generated by the quantal conductance represented as a multiexponential function (Equation (1)) with a peak amplitude of −12.783 nS, τ1 = 0.5 ms, and τ2 = 3 ms [7].

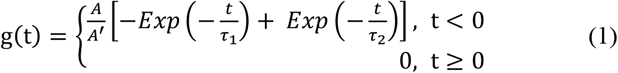

where g(t), τ_1_, and τ_2_ are the quantal conductance, rise, and decay time constant, respectively (τ_2_ > τ_1_). A is the peak amplitude, and A’ is a normalization factor:

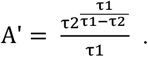

To generate variation in PSC events, a random number drawn from a normal distribution with mean of 1 and standard deviation of 0.3 was multiplied with time constants, τ_1_ and τ_2_. A total of 250 simulated PSCs were generated and positioned randomly in a 250-sec long idealized waveform. To test effects of the background noise on event detection performance for the package, three types of background noise with standard deviation (σ_noise_) of 2, 6, and 10 pA were added to the idealized waveform with simulated events, respectively. Referring to the position of simulated PSCs in the idealized waveform, we computed a confusion matrix containing the number of true positive (TP), true negative (TN), and false positive (FP), and false negative (FN) to summarize the detection performance of the package under the effects of three background noises, respectively. Based on the confusion matrix, the precision and recall were defined as follows: precision = TP/(TP+FP), recall = TP/(TP+FN). The F1 score is the harmonic mean of precision and recall (2*(precision*recall)/(precision+recall)). All procedures of simulation were performed using R Project for Statistical Computing (RRID:SCR_001905).

### 2.3 Experimental recording of PSCs

#### 2.3.1 Animal

C57BL6/J male mice aged 7-week-old were used. Mice were housed with food and water available *ad libitum* under a 12 hours light/dark cycle. All animal use was in accordance with protocols approved by the Animal Care and Use Committee of Seoul National University College of Medicine.

#### 2.3.2 Slice preparation

Brain slices were prepared as described previously [11]. Brain slices (250 μm thick) were obtained using a vibratome (VT1200, Leica) after isoflurane anesthesia. Slices were cut in a chamber filled with ice-cold cutting solution, NMDG-HEPES, composed of the following (in mM): 2.5 KCl, 1.25 NaH_2_PO_4_, 93 NMDG, 30 NaHCO_3_, 20 HEPES, 25 glucose, 5 sodium ascorbate, 2 Thiourea, 3 sodium pyruvate, 12 L-acetyl-cysteine, 10 MgSO_4_•7H_2_O and CaCl_2_•2H_2_O bubbled with 95% O2 and 5% CO2. The slices were immediately put into an artificial CSF (aCSF) composed of the following (in mM): 125 NaCl, 2.5 KCl, 1 MgCl_2_, 2 CaCl_2_, 1.25 NaH_2_PO_4_, 26 NaHCO_3_ and 10 glucose bubbled with 95% O_2_ and 5% CO_2_.

#### 2.3.3 sIPSC Recording in the prelimbic pyramidal neurons

Slices were transferred to a recording chamber perfused with oxygenated aCSF at 30-32°C controlled by a peristaltic pump. Cells in the prelimbic region were visualized with 40x magnification objective (Olympus) on the stage of upright microscope (BX61W1, Olympus) equipped with infrared-differential interference contrast optics in combination with digital camera (AquaCAM Pro/S3). Patch microelectrodes were pulled from borosilicate glass (O.D.:1.5mm, I.D.: 1.10mm, WPI) on a Flaming-Brown micropipette puller model P-1000 (Sutter Instruments, USA). Patch microelectrodes had a resistance of 4-8MΩ. Signals were recorded using a patch-clamp amplifier (Multiclamp 700B, Axon Instruments, USA) and digitized with Digidata 1550A (Axon Instruments, USA) using Clampex software. Signals were amplified, sampled at 10 kHz, and filtered to 2 or 5 kHz. Pyramidal neurons were identified by large apical dendrites. During voltage-clamp recordings, spontaneous inhibitory postsynaptic current (sIPSC) was recorded at −60mV membrane holding potential with high chloride intracellular solution (in mM): 150 CsCl, 2 MgCl_2_6H_2_O, 0.1 CaCl_2_2H_2_O 10 HEPES, 1 EGTA, 2 Na-ATP, 0.4 Na-GTP respectively. During sIPSC recording, 10 μM NBQX (2,3-dihydroxy-6-nitro-7-sulfamoyl-benzo[f]quinoxaline) and 50 μM AP-5 (Tocris Bioscience, UK) were applied in the bath to block excitatory synaptic responses.

Recorded sIPSC was analyzed using two programs employing different detection algorithms, MiniAnalysis (Synaptosoft, USA) and Clampfit (Molecular Devices, USA), respectively, as well as Minhee Analysis Package. As for MiniAnalysis, the amplitude and area thresholds were set to 10 and 6.9462, respectively. In regard with Clampfit, the event template was created according to the user manual. Template match threshold was adjusted to 6. A Kolmogorov-Smirnov test was used to compare the pairs of results from three different detection programs. The level of statistical significance was set to p ≤ 0.05.

#### 2.3.4 sEPSC recording in the cerebellar Purkinje cells

Slices were placed in a submerged chamber on the stage of a microscope (BX50WI, Olympus Optical, Japan) and perfused with aCSF. The whole-cell voltage-clamp recordings were performed from PCs in the cerebellum at 32°C using the recording patch pipettes (2.5-3.5 MΩ) filled with internal solution containing the following (in mM): 140 Cs-methanesulfonate 4 NaCl, 0.5 CaCl_2_, 10 HEPES, 2 MgATP, and 5 EGTA, pH 7.3 accompanied with Multiclamp 700B (Molecular Devices) and Digidata 1440A (Molecular Devices). The sampling frequency of 10 kHz and filtering of signals at 2 kHz was kept constant throughout the experiment. All of the recordings were conducted within the aCSF containing 100 μM picrotoxin (Sigma-Aldrich, USA) to block inhibitory synaptic inputs onto the Purkinje cells. All sEPSC recordings were acquired in the Lobule Ⅲ-Ⅴ or Lobule Ⅹ of cerebellar vermis.

## 3. RESULTS

### 3.1. Package overview

The Minhee Analysis Package is a standalone program that operates under Windows platform using Microsoft Windows 7 or later. For the present study, the package was installed and run on a general 64-bit desktop computer with an intel^R^ core™ i3-2100 CPU and 8GB RAM. Minhee Analysis Package includes two independent programs, Minhee Analysis and Retriever (Figure 1). The main features of each program are detailed in the USE EXAMPLES section below. The package provides an integrated GUI that allows users to execute all functions and visualize the results. The GUIs of the package can be minimized, but not resizable, therefore, the package requires a computer monitor with at least 1336*768 resolution.

**Figure 1.**
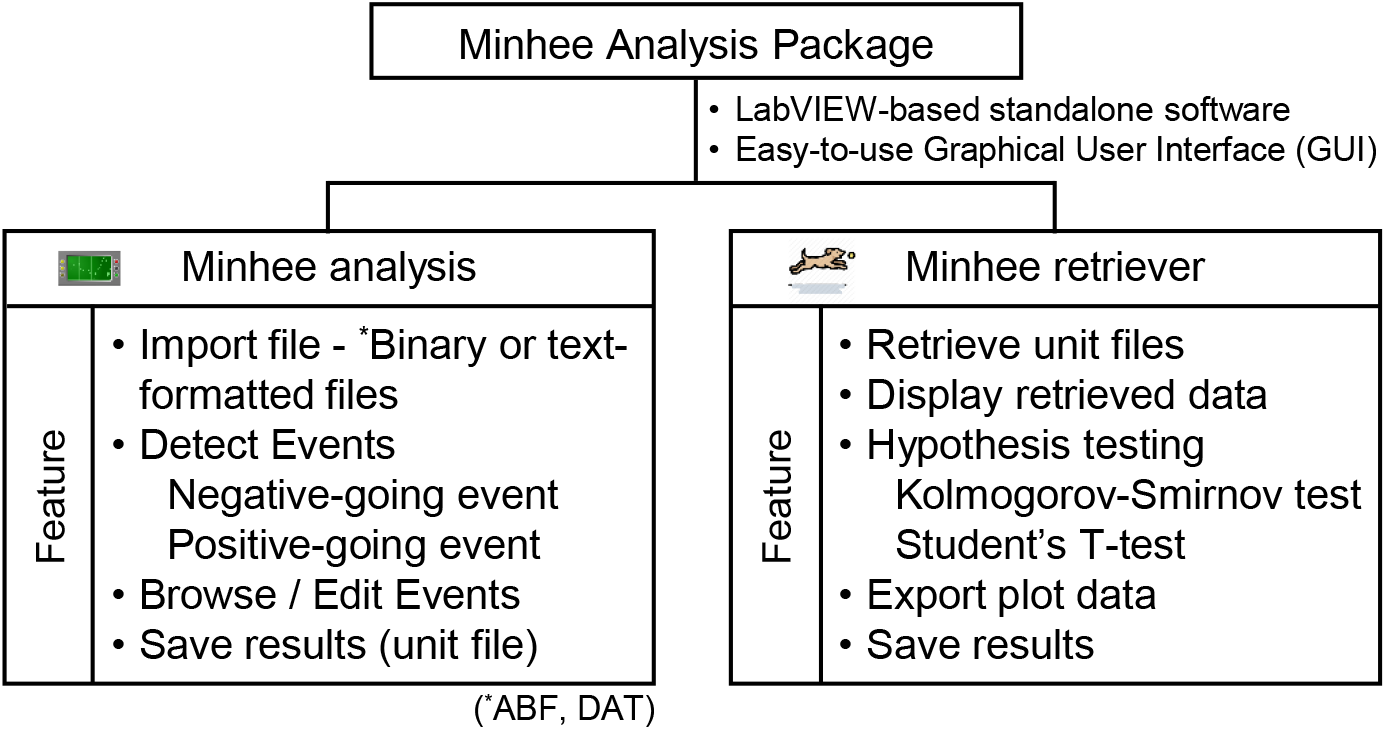
Package overview. The Minhee Analysis Package consists of two independent applications: Minhee Analysis and Retriever. This package is a standalone software developed by the LabVIEW (National Instrument, USA). Each of the two applications provides an easy-to-use graphical user interface (GUI) that allows a user to execute all the functionality and displays the result. The main features of each application listed at the bottom of the chart are described in the use examples.

### 3.2. Algorithm for event detection

To detect PSC events, the Minhee Analysis Package utilizes a step-by-step search algorithm using user-defined and built-in functions from the LabVIEW development environment. The detection procedure includes four steps: smoothing, initial search, baseline search, and final search (Figure 2). First, the smoothing is applied to remove high-frequency noise in raw data (Figure 2a and b). In the second step, the peak (or valley) of signals with the amplitude exceeding 3 pA from the trend of the recorded trace is accepted as local peaks (or valleys) (Figure 2c). Once the location of the local peak (or valley) is defined, the algorithm begins a backward search for the baseline of the local peak (or valley) (Figure 2d). In this step, the algorithm uses the second derivative of the trace to determine the local minimum (or maximum) and then considers defined local minimum (or maximum) to be the baseline of the event. In the final step, the algorithm calculates the amplitude of the event as the difference between the peak (or valley) and baseline, and then finally determines the events (Figure 2e). Only events with the amplitude exceeding over the user-defined criterion are accepted. Otherwise, the events are discarded.

**Figure 2.**
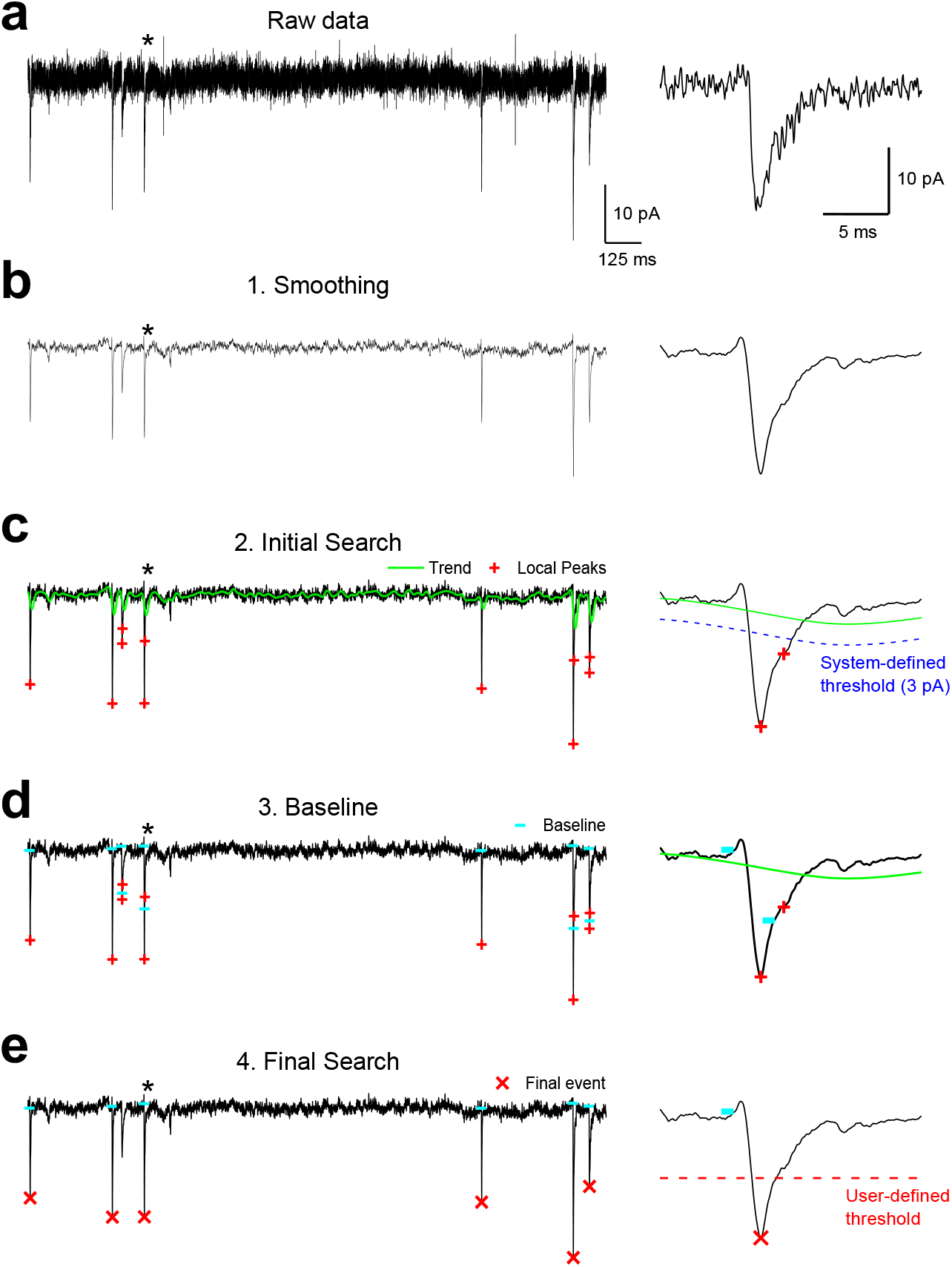
The scheme of event detection. The process of event detection consists of four steps. The asterisk represents the region of the event shown at an extended timescale in the right panel of each row. (a and b) Raw traces were smoothed using an S-G filter (c) Any valley point below the amplitude threshold (3 pA, dashed blue line) considered as a local minimum (red plus symbol). (d) The baseline of a local minimum was found using a backward search algorithm (cyan horizontal bar). (e) The final event (red x symbol) was determined by comparing the amplitude of the event with the minimum amplitude that a user established (dashed red line). In (c and d), the green line indicates the trend of trace.

### 3.3. Validation of the event detection algorithm using simulated data

We first tested the validity of event detection algorithm using simulated data (Figure 3). EPSCs were simulated with variable amplitude and kinetics (Figure 3a), and they were inserted in an idealized waveform with random interevent interval (IEI) (Figure 3b). With 250 EPSCs generated, the averaged amplitude of the total simulated EPSCs was 27.85 ± 2.19 pA (mean ± S.D.). The rise and decay time constants were 0.299 ± 0.059 ms and 3.04 ± 0.057 pA, respectively. To evaluate the algorithm, three noise-added waveforms were analyzed by Minhee Analysis (Figure 3c). Despite the increase of the background noise in the waveforms, the program accurately detected simulated EPSCs (Figure 3d). As expected, event detection results in low (σ_noise_: 2 pA), moderate (σ_noise_: 6 pA), and high (σ_noise_: 10 pA) noise conditions exhibited a similar distribution of events with that of the original EPSCs. In σ_noise_ = 10 pA, the algorithm missed only 0.4% of the events and detected five false-positive events (26.42 ± 3.86 (mean ± S.D.)).

**Figure 3.**
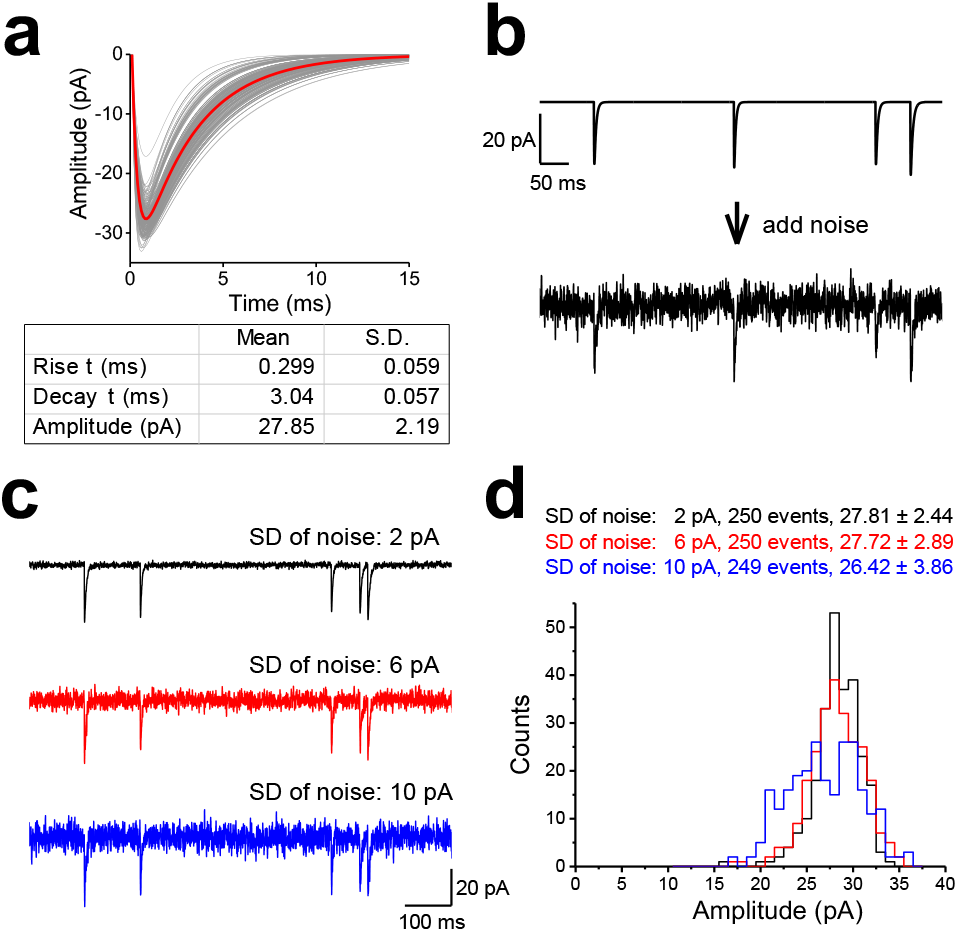
Validation of the event detection algorithms using simulated data. (a) Artificial EPSCs were randomly generated by the multiexponential equation (eq. 1). To generate various kinetics of EPSCs, the rise and decay time constants (τ) were randomly scaled by multiplying them with a random factor (mean 0, standard deviation 0.3). The individual and averaged traces of the simulated EPSCs are in gray and red, respectively. (b) White noise was generated according to the normal distribution, and then added to the idealized waveform (top, idealized trace; bottom, noise-added trace). (c) Representative traces of the simulated EPSC generated with three levels of background noise. The standard deviations (SD) of noise were 2 pA (top, black), 6 pA (middle, red), and 10 (bottom, blue), respectively. (d) A histogram showing the amplitude of the EPSC detected in the simulated traces. For SD of noise = 2 and 6 pA, the adjustable parameters were as follows: *Minimal amplitude* = 10, *Polynomial Order* = 3, *Side Points* = 14. For SD of noise = 10 pA, while the *Polynomial Order* was the same as others, the *Minimal Amplitude* and *Side Points* were adjusted to 15 and 20, respectively, to minimize the number of false-positive events.

### 3.4. Demonstration of how parameter adjustments affect event detection

In the Minhee Analysis, we implemented a step-by-step search algorithm employing a variety of functions in the LabVIEW development environment. The performance of event detection depends on several adjustable parameters of the algorithm. These parameters should be easily understood by a user and the number should be as small as possible. In the Minhee Analysis, we simplified the types of parameters that a user needs to adjust as parameters related to the smoothing step (***Smoothing parameters***) and parameters related to the shape of the event (***Event detection parameters***). In the smoothing step of the algorithm, ***Polynomial order*** and ***Side points*** are the adjustable parameters, which have a conflicting effect on the smoothness. A higher ***Polynomial order*** leads to a less smoothed signal while a higher ***Side points*** yields a more smoothed signal (Figure 4a and b). In addition to the Smoothing parameters, the ***Minimum amplitude*** also should be defined by users in the final search step. Optimizing these parameters is required for successful event detection, which not only minimizes false-positive cases but also maximizes true-positive cases. Therefore, we demonstrated the effect of adjustable parameters on the accuracy of event detection as each parameter was independently manipulated.

**Figure 4.**
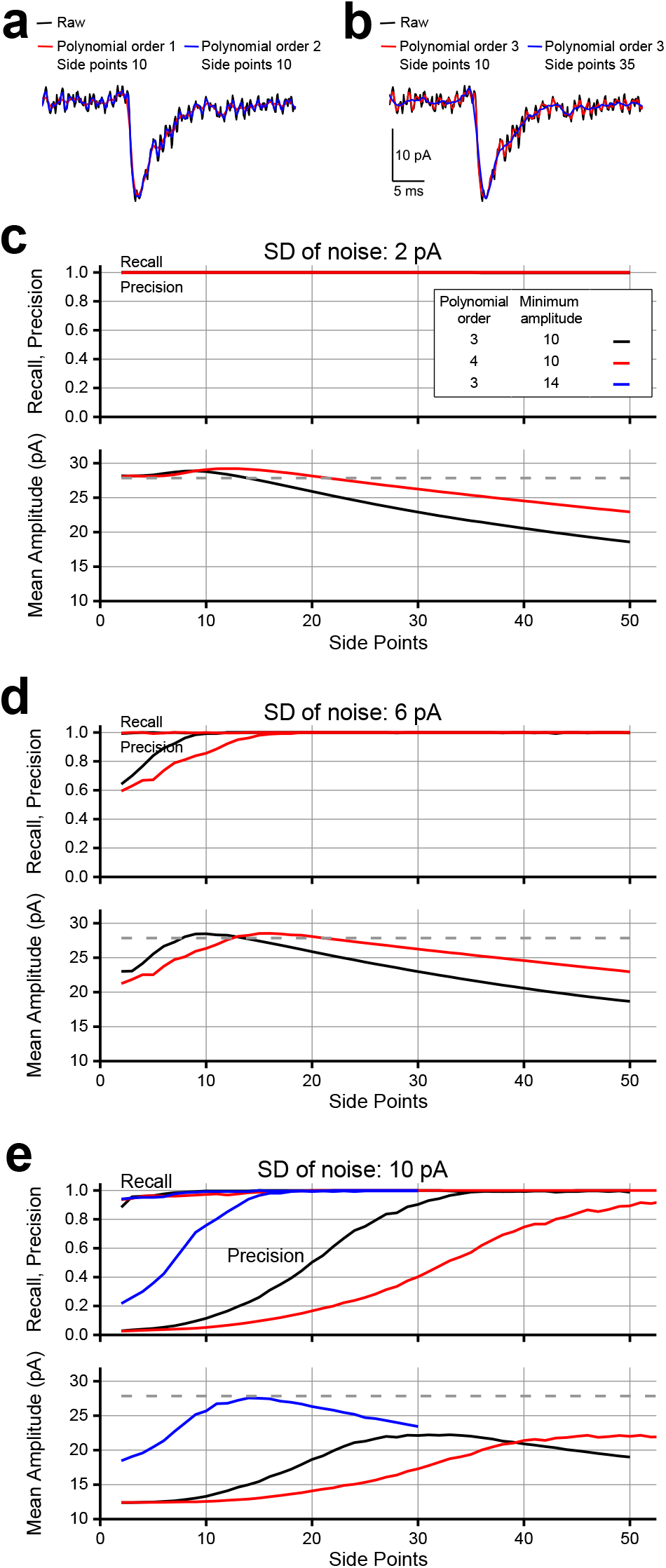
Demonstration of adjustable parameters for successful event detection. Effects of smoothing parameters, Polynomial order (a) and Side points (b) on the smoothness of the traces. The recall, precision (top), and mean amplitude (bottom) of the events were calculated from simulated data with SD of noise: 2 pA (c), 6 pA (d), and 10 pA (e). In (c-e), the dashed line indicates the mean amplitude of the original simulated EPSCs.

First, we performed event detection on three noise-added simulated data under the range of ***Side points*** from 2 to 50, and ***Polynomial order*** of 3 or 4, while ***Minimum amplitude*** was fixed to 10. Next, we generated a confusion matrix containing the number of TP, TN, FP, and FN cases for each result of event detection. Finally, to assess the accuracy of the algorithm, F1 scores were calculated for a wide range of ***Side points*** values, which consider both the recall and precision of the detection (see Materials and Methods). Additionally, since a number of FP or excessive removal of noise could underestimate the mean amplitude of detected event without intention, we summarized the mean amplitude of detected events over the range of smoothness to compare with the mean amplitude of the simulated events.

In the result of σ_noise_ = 2 pA, our algorithm accurately detected events over the range of parameters that we tested (Figure 4c). The mean amplitude of detected events exhibited a unimodal pattern against the range of ***Side points*** from 2 to 50. In ***Polynomial order*** of 4, the peak of the mean amplitude shifted to the right compared to the result in ***Polynomial order*** of 3, and the graph decreased gently, indicating that adjusting ***Side points*** at a high ***Polynomial order*** has a less robust effect on the smoothness of the traces than adjusting ***Side points*** at a low ***Polynomial order***. As for the result of σ_noise_ = 6 pA, when ***Polynomial order*** was set to 3, events were reliably detected at a ***Side points*** equal to or greater than 7 (F1 score > 0.95) (Figure 4d). The minimum ***Side points*** was 12 to exceed F1 score of 0.95 when ***Polynomial order*** was set to 4. Considering each result of the mean amplitude, ***Side points*** 7 to 16 and 12 to 26 were in appropriate ranges to maximize the accuracy of event detection in ***Polynomial order*** of 3 and 4, respectively. As expected, increasing ***Side points*** was necessary to achieve a F1 score over 0.95 in the high noise condition (σ_noise_ = 10 pA) (Figure 4e). In regard with the mean amplitude of the high noise condition, the maximum mean amplitudes of events reduced 20% (***Polynomial order*** 3 and ***Side points*** 33, 22.23 pA; ***Polynomial order*** 4 and ***Side points*** 51, 22.15 pA) as a result of smoothing. Lastly, when ***Minimum amplitude*** was set to 14, the mean amplitude of events was restored to 99% of the mean amplitude of the raw simulated events.

In summary, we demonstrated the effect of the adjustable parameters on the accuracy of the algorithm using the noise-added simulated EPSCs. For a successful event detection, these parameters should be optimized within a specific range in response to noise levels.

### 3.5. Comparison of different event detection methods

Next, the performance of our algorithm using experimental data was compared with the detection by other programs employing different algorithms. Experimental data were prepared by recording spontaneous inhibitory postsynaptic current (IPSC) in a pyramidal neuron of the medial prefrontal cortex in a mouse. Spontaneous IPSCs were detected by two commonly used analysis tools: MiniAnalysis (Synaptosoft, USA) [1, 2, 12–14], and Clampfit (Axon Instruments, USA) [15, 16].

The numbers of events detected by Minhee Analysis and MiniAnalysis were 173 and 177, respectively. By Clampfit, we initially detected 168 events, but discarded 23 events that exhibited amplitudes less than 10 pA (Table 1 and Figure 5). Despite the minor discrepancy across the results, the quantification of total events by Minhee Analysis was comparable with those by others (amplitude: Minhee Analysis, 19.144 ± 1.8 pA; MiniAnalysis, 19.205 ± 1.872 pA; Clampfit, 20.733 ± 2.053 pA; inter-event interval: Minhee Analysis, 1.399 ± 0.232; MiniAnalysis, 1.388 ± 0.213; Clampfit, 1.605 ± 0.274). No significant differences were observed in comparisons of the amplitude of events and inter-event interval (p > 0.05, amplitude; p > 0.05 inter-event interval; Kolmogorov-Smirnov test). In these comparisons using experimental data, we confirmed the fidelity of our event detection algorithm.

**Table 1.**
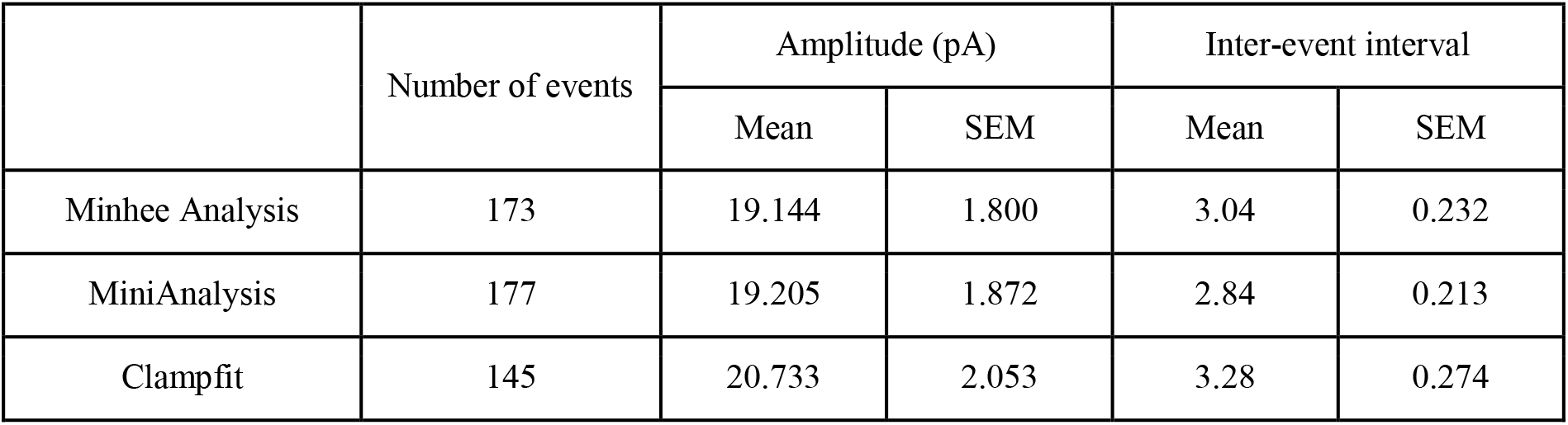
The number and descriptive statistics of total events made by Minhee Analysis, MiniAnalysis (Synaptosoft, USA), and Clampfit (Axon instrument, USA).

|  | Number of events | Amplitude (pA) |  | Inter-event interval |  |
| --- | --- | --- | --- | --- | --- |
|  |  | Mean | SEM | Mean | SEM |
| Minhee Analysis | 173 | 19.144 | 1.800 | 3.04 | 0.232 |
| MiniAnalysis | 177 | 19.205 | 1.872 | 2.84 | 0.213 |
| Clampfit | 145 | 20.733 | 2.053 | 3.28 | 0.274 |

**Figure 5.**
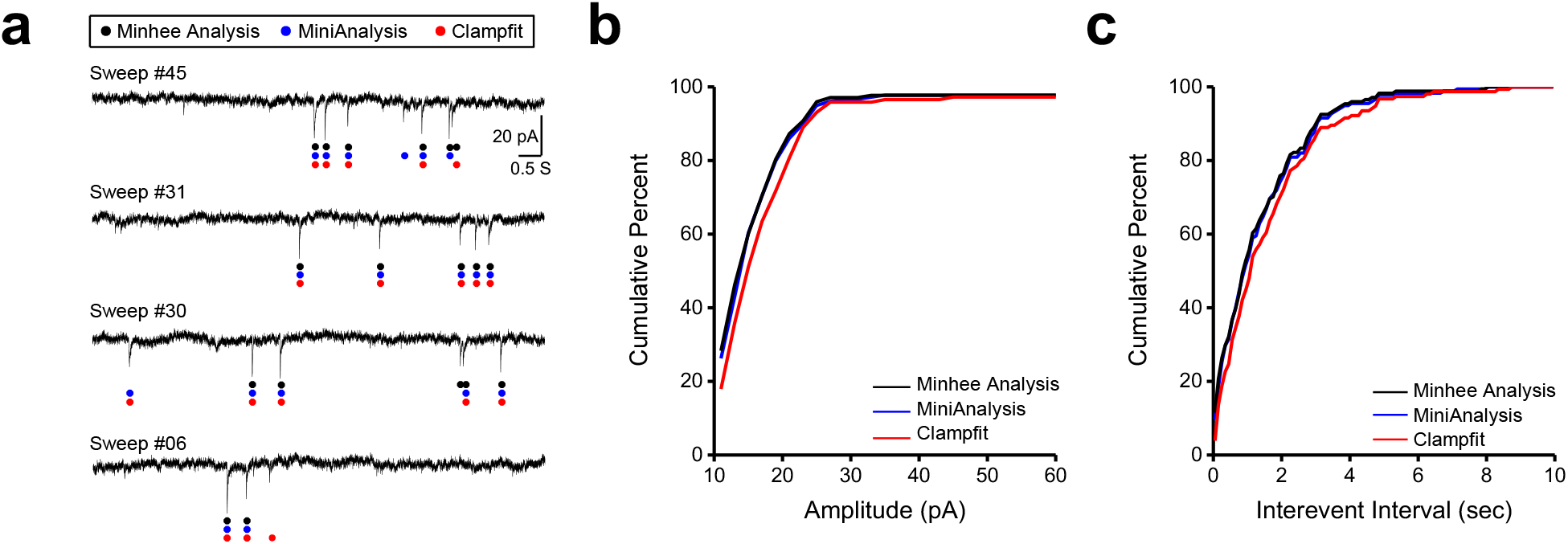
Comparison of different detection methods. (a) Representative traces of sEPSCs with results of detecting event by different methods. Red, blue, and green dots denote events detected by Minhee Analysis, MiniAnalysis, and Clampfit, respectively. (b) Cumulative distribution of the amplitude. (c) Cumulative distribution of IEI. The results of Minhee Analysis, MiniAnalysis, and Clampfit are depicted in black, blue, and red lines, respectively, in (b) and (c).

## 4. USE EXAMPLES

Here, we describe examples of analysis and management of spontaneous synaptic events using Minhee Analysis (Figure 6) and Retriever (Figure 7). These examples are intended to describe the functionality of the package and general workflow for a successful analysis using the package. The package includes all the data used in these examples in the Example Files folder (C:\Program Files (x86)\NeuroPhysiology Lab\Example Files), therefore, a user can reproduce the results in these examples. In this section, the name of the GUI and the function of the package are shown in bold.

**Figure 6.**
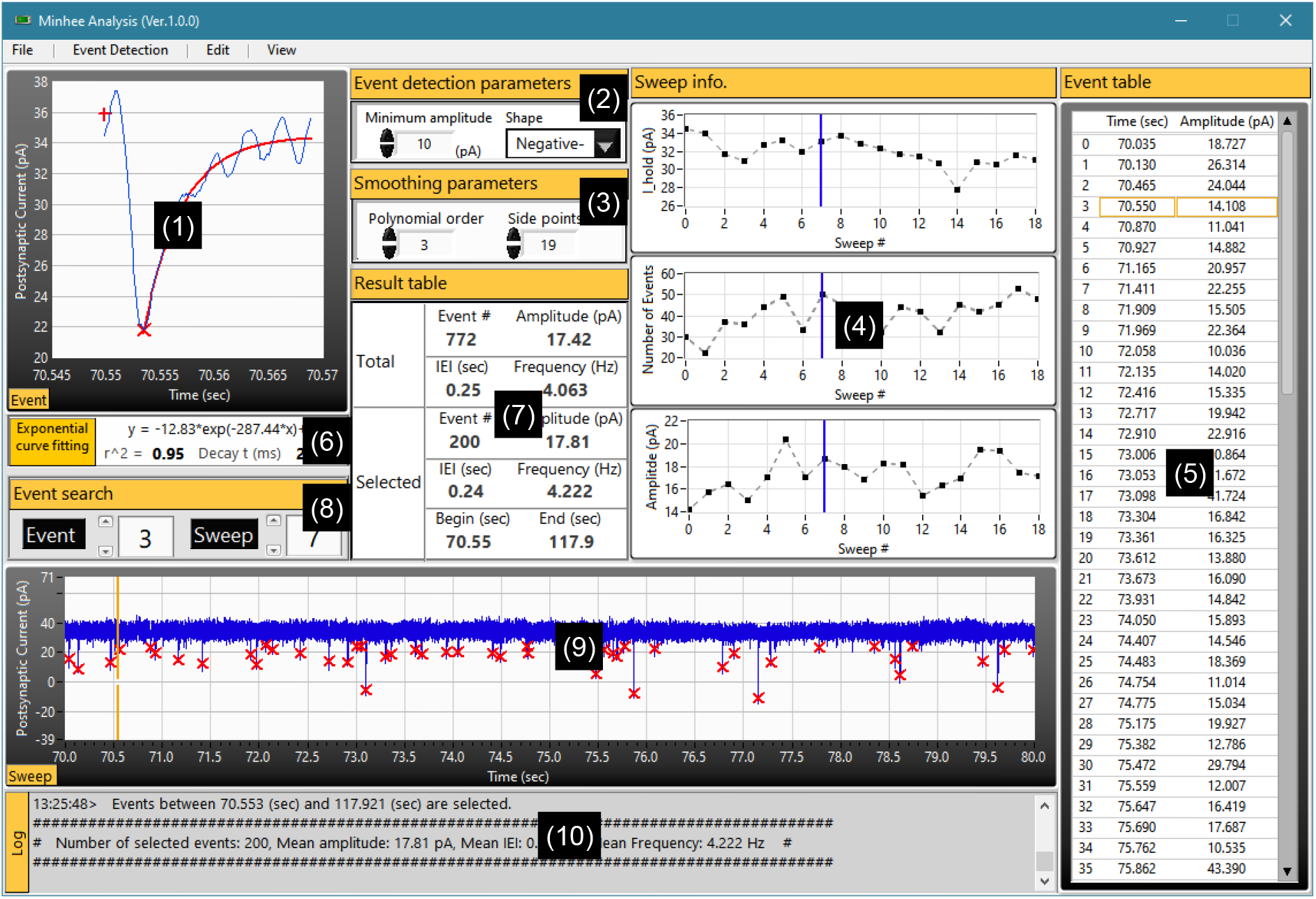
Screen capture of Minhee Analysis’s GUI. The GUI of the program has 10 subsections titled with section names in yellow boxes (numbers in parentheses). (1) The Event window displaying the selected event in the Event search panel. The baseline, peak, and exponential curve fits are displayed as the plus, multiply sign, and red line, respectively. (2) Event detection parameters. (3) Smoothing parameters. (4) Multiple graphical windows showing the overall trend of the data by sweeps. The top, middle, and bottom plots represent averaged holding current, the total number of events, and averaged amplitude of events in a sweep, respectively. Blue vertical lines indicate the selected sweep in the Event search panel. (5) Event table exhibiting the time and amplitude of events from a selected sweep located in the Event search panel. The orange boxes indicate the selected event in the Event search panel. (6) Exponential Curve Fitting panel (7) Summary of the detected events. The Total section shows the total number of events, averaged amplitude, and IEI from the whole event. In the Selected section, those are calculated only from the events belonging to the distribution of events that a user made. (8) Event search specifying the index number of events or sweep to display in Minhee Analysis’s plot windows. (9) The Sweep window displaying the whole trace of the selected sweep. The orange vertical bar indicates the selected event in the Event search panel. (10) Log window displaying the history and results of executed commands.

**Figure 7.**
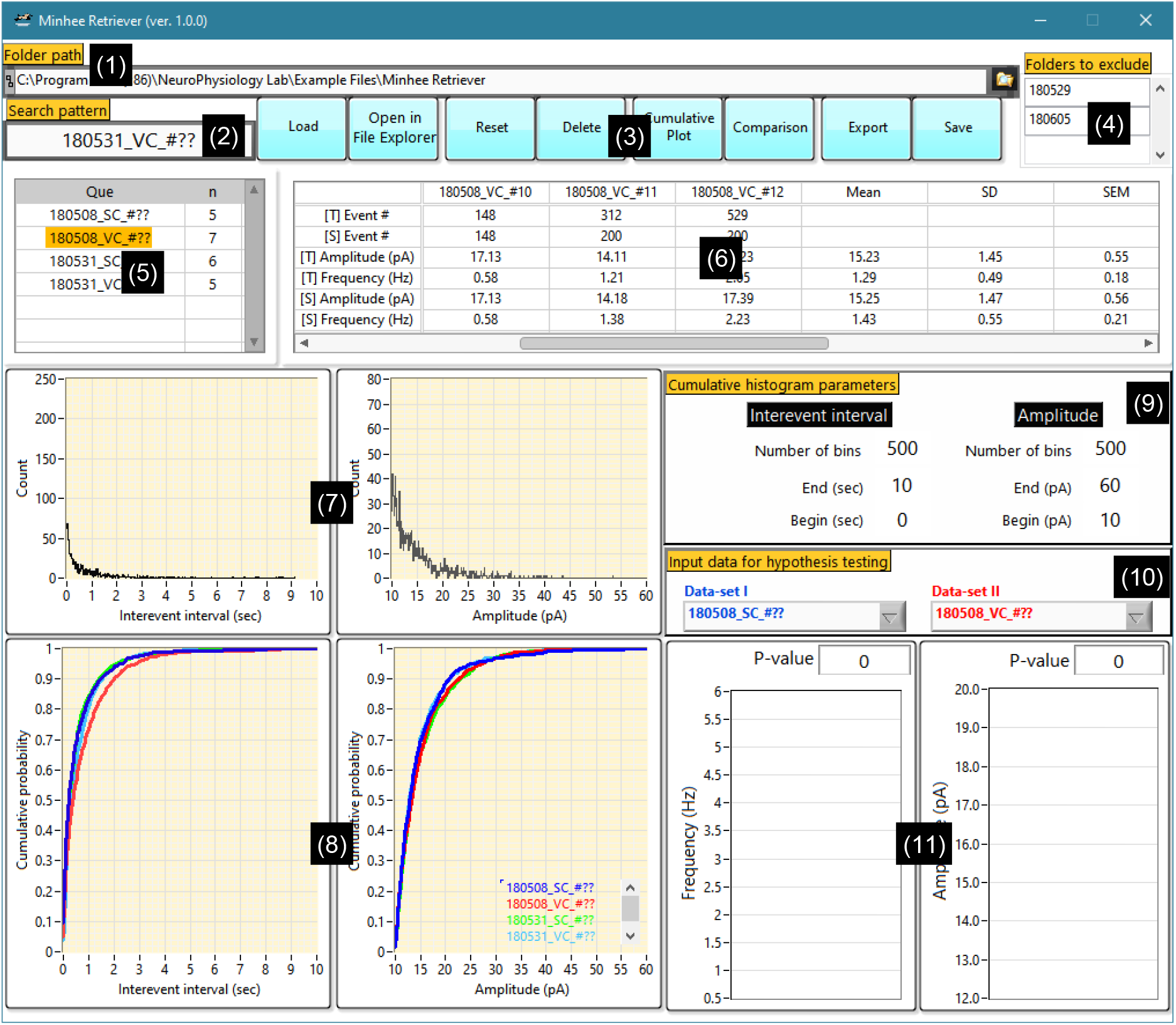
Screen capture of Minhee Retriever’s GUI. The GUI of the program has 11 subsections (numbers in parentheses). (1) The Folder Path is used to specify the top-level folder that contains the unit file. (2) A user can enter search patterns to retrieve unit files. (3) Several Action buttons are available to execute the functions. (4) The unit files listed here are excluded the during retrieval. (5) Query table. This table shows currently retrieved results. (6) The result table shows the number of events, amplitude, and frequency of the total and selected events in each unit file. General (7) and cumulative relative (8) histograms for the IEI and amplitude of selected events. Parameters for generating the cumulative relative histogram. (10) For statistical comparison between two groups, two data-sets are determined by a user. Result of hypothesis testing is displayed in (11).

### 4.1. Installing Minhee Analysis Package

1. Download all files from the Volume folder in the following github repository (https://github.com/parkgilbong/Minhee_Analysis_Pack)
2. Run setup.exe
3. After the installation, two executable files, Minhee Analysis.exe and Minhee Retriever.exe, can be found in the folder located in C:\Program Files (x86)\NeuroPhysiology Lab.

### 4.2. Use example of Minhee Analysis

#### 4.2.1. Import file and Browse traces

Minhee Analysis accepts binary (ABF and DAT) and text-formatted (TXT) files (Table 2). The ABF and DAT format are the most common file types in the electrophysiological field, which is created by pClamp software (Molecular Devices, USA) and PATCHMASTER (HEKA Elektronik, Germany), respectively. An ABF file (ver 1.8 or higher) created under the episodic simulation mode can be directly readable by Minhee Analysis. It is noted that the PATCHMASTER uses a data tree of five levels (Root-Group-Series-Sweep-Trace) to manage its data, therefore, a DAT file (v2) stored in the data tree as series with multiple sweeps can be imported into the package. Each sweep should contain a PSC Trace. To import a DAT file, a user should select one item at the Series level (Figure 8a).

**Table 2.** List of files used and created by Minhee Analysis Package

| File type | Description |
| --- | --- |
| ABF | Axon binary file created by pClamp 9 or earlier (ABF 1.83 or 1.84)<br>Axon binary file created by pClamp 10 or later (ABF 2.0)<br>Data acquisition mode: Episodic stimulation mode.<br>Recommended sampling rate: 10 – 20 kHz<br>Read-only in Minhee Analysis |
| DAT | Binary file generated by PATCHMASTER<br>A bundle file with merged the Raw Data File (*.dat) and Acquisition Parameter File (*.pgf)<br>Multi-sweep data stored in the Series level of the data tree<br>Recommended sampling rate: 10 – 20 kHz<br>Read-only in Minhee Analysis |
| TXT<sup>a</sup> | Tab-delimited without header<br>Two-dimensional array with the sweeps organized column-wise<br>Data unit: A or pA<br>Recommended sampling rate: 10 – 20 kHz<br>Read-only in Minhee Analysis<br>Additional parameters related to file properties, such as sampling rate and sweep duration, are required as a file import. |
| TXT<sup>b</sup> | Three rows delimited by tabs without headers (1st column, time of event (sec); 2nd column, amplitude of event (pA); 3rd column, selection index)<br>Write-only in Minhee Analysis and read-only in Minhee Retriever |

<sup>a</sup> Experimental recording data exported from a third-party software
<sup>b</sup> Event file created by Minhee Analysis (referred as the unit file throughout the current article)

**Figure 8.**
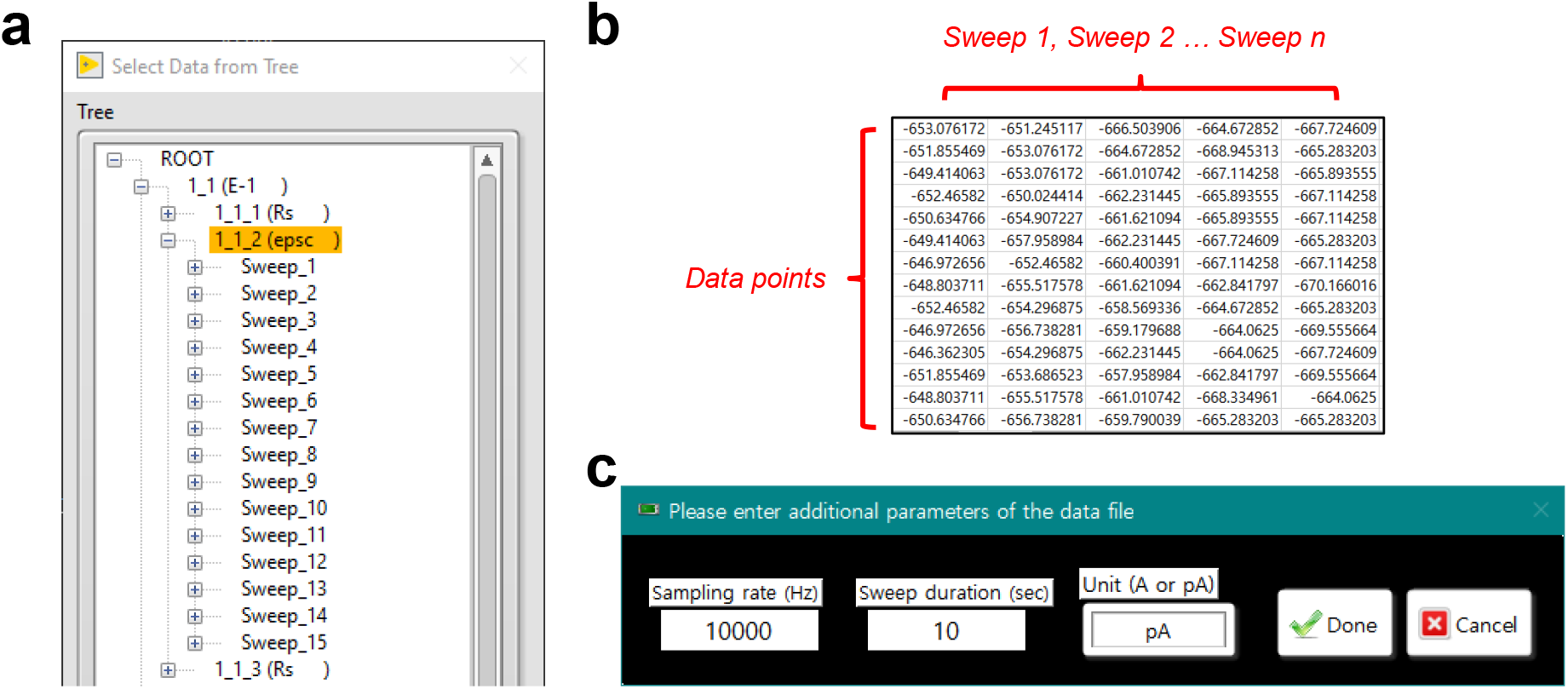
Importing data files into Minhee Analysis. (a) To import a DAT file generated by the PATCHMASTER, a user should select one item at the series level in the data tree. (b) Structure of text-formatted data file. (c) A pop-up window for entering additional parameters when importing a text-formatted file.

In addition to the two binary files, a TXT file of experimental data can be read by Minhee Analysis. In this format, data of each sweep organized in columns and a unit of data stored in a TXT file must be A or pA (Figure 8b). As a user is importing a file, additional parameters related to experimental recording, such as sampling rate, sweep duration, and unit of stored data, should be specified (Figure 8c). For a proper recording, the sampling frequency of 10 - 20 kHz is recommended. A file with the sampling frequency over 20 kHz or recorded within one very long sweep under the gap-free mode may cause shutdown of the package due to insufficient memory.

Once a file is imported in the Minhee Analysis, users are able to browse traces by using the navigation keys in ***Event search*** (Figure 6, (8)). The selected sweep trace is displayed in ***Sweep viewer*** (Figure 6, (9)). The numbering system for the package is zero-based, which the first sequence of elements starts from index zero. Initial inspection of the data can be done in this step before the actual event detection.

#### 4.2.2. Set parameters and Detect events

There are four adjustable parameters in the Minhee Analysis (Figure 6, (2) and (3)). First, users can choose the shape of events between negative-going and positive-going events (Figure 9). The shape of events is determined by electrophysiological properties, such as holding potential and ionic concentration in an intracellular solution. Then, users can specify ***Minimum amplitude***, a criterion for judging valid events. The default ***Minimum amplitude*** is set to 10 (pA). Lastly, users can adjust ***Polynomial order*** and ***Side points***. As we demonstrated in the results (Figure 4), increasing these two parameters has the opposite effect on the smoothness of signal. Therefore, it is recommended that users change only one parameter, while the other is fixed, for the analysis.

**Figure 9.**
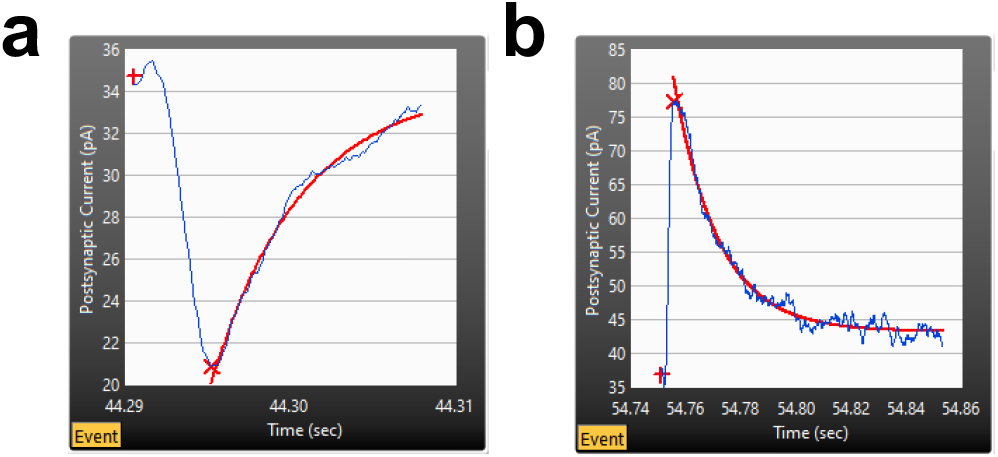
Negative- (a) and positive-going (b) events can be detected by Minhee Analysis. In each panel, the baseline, (negative or positive) peak, and exponential curve fits are displayed as a plus symbol, multiplication sign, and red line, respectively

After optimization of four parameters, a user can execute ***Detection*** (*Event Detection>Detection*) for searching PSC events. While event detection is in progress, the results are updated in real-time in ***Sweep info*** (Figure 6, (4)). In ***Sweep info***, the averaged holding potential, number of detected events, and mean amplitude of the detected events from each sweep are displayed on the top, middle, and bottom graphs, respectively. The Minhee Analysis also supports ***Abort*** (*Event Detection>Abort*) function to disrupt the process.

#### 4.2.3. Browse the result

Once the detection is completed, the results are displayed on each panel of the Minhee Analysis (Figure 6). The displayed information on the panels of the Minhee Analysis is divided into two categories. One is information related with individual events, and the other is information about each sweep. To display a sweep trace, users can use the navigation keys or manually enter the index number of the sweep of interest. Each chart in ***Sweep info*** has a blue vertical cursor indicating the currently selected sweep. Users can refer to ***Sweep info*** to see trends in the data across the sweep.

The peak time and amplitude of whole events in a sweep come up in ***Event table*** (Figure 6, (5)). Users can browse an event trace in ***Event viewer*** (Figure 6, (1)). The index number of the selected event in ***Event search*** synchronizes with the yellow boxes and cursor in ***Event table*** and ***Sweep viewer***. ***Exponential curve fitting*** panel shows the result of curve fitting using the following single exponential function with the data after the peak point of each event (Figure 6, (6)). A user can refer to the result of curve fit to determine whether false positive events exist.

#### 4.2.4. Edit events

False-positive or -negative events can be included in the result of event detection. A user is able to edit the detected events using ***Redetect*** and ***Delete*** functions (*Edit>Redetect, Delete*). Redetect is a function of redoing the process within the selected sweep, and it is useful to find a false-negative event. For this purpose, a user can adjust the event detection parameters that apply only to the currently selected sweep. An example of use is shown in Figure 10a. While the rest of the events remained the same, one false-negative event was detected by switching ***Side points*** from 25 to 27. ***Delete*** function can be used to eliminate any false-positive event (Figure 10b). A user can distinguish a false-positive event among actual events based on its shape and curve-fitting of the event. On the other hand, if a true-positive event is deleted, it can be simply restored by ***Redetect*** function.

**Figure 10.**
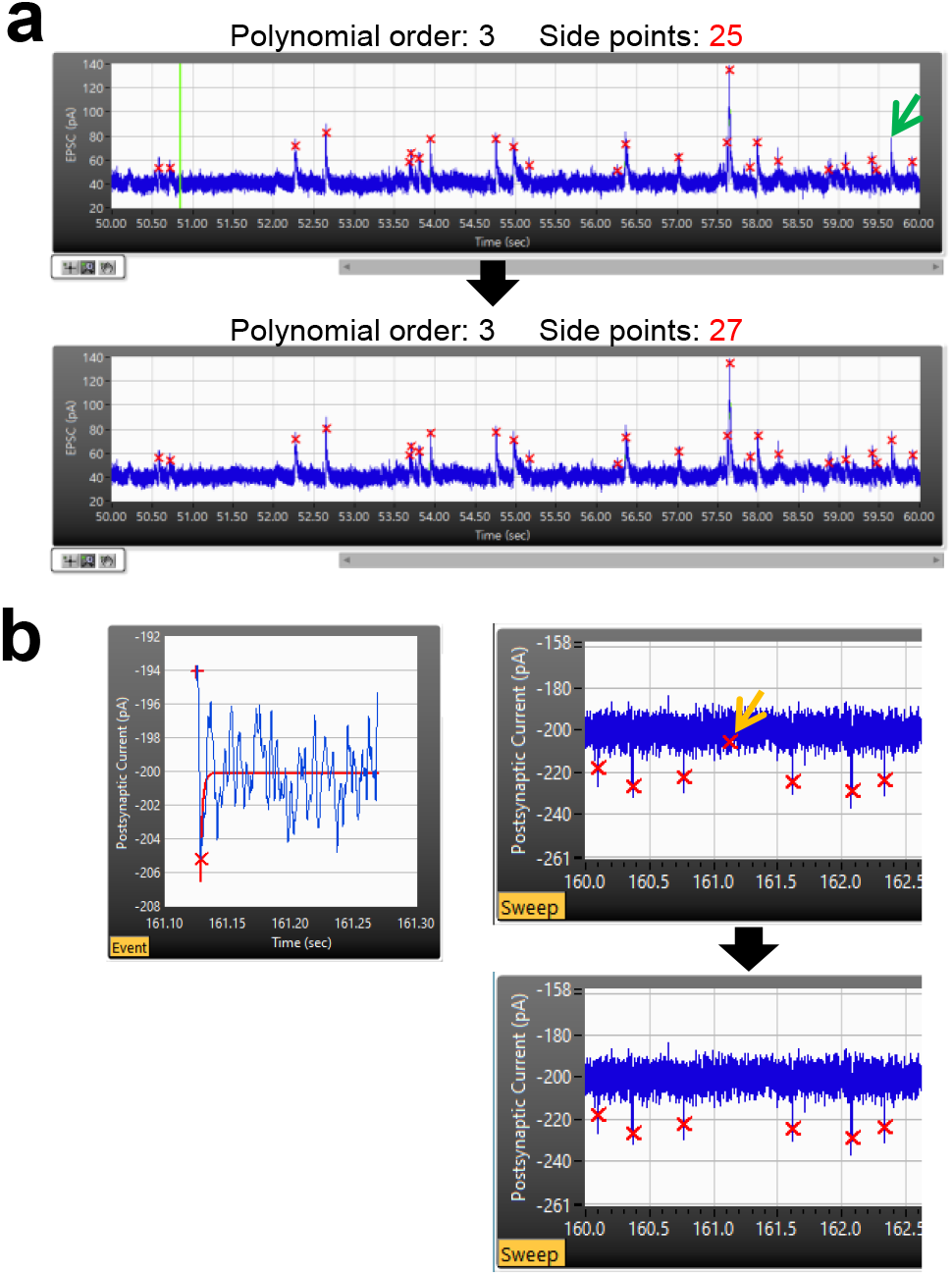
Editing events in Minhee Analysis. (a) Detection of a false-negative event (green arrow) by executing Redetect function with optimal parameters (bottom). (b) By executing the Delete function, a false-positive event (left and yellow arrow in the top right) was excluded.

#### 4.2.5. Select events

Spontaneous synaptic events of a neuron exhibit a broad range of properties in terms of measurements such as the IEI and amplitude. Therefore, a sufficient number of events are required to characterize the synaptic properties of a single neuron. To this end, Minhee analysis has ***Select*** (*Edit>Select*) function. Select function allows a user to select as many sequential events as desired. Before executing ***Select*** function, users specify the desired number of events in the pop-up window (Figure 11a) and update ***Event search*** parameters to choose the beginning of the sequential events. ***Sweep info*** helps a user to estimate whether events have been selected under stable conditions. After ***Select*** function executes, results calculated only from the events belonging to the distribution of events created by a user will be displayed in the ***Selected*** section of ***Result table*** (Figure 6, (7)). A user can compare the results between the ***Total*** and ***Selected*** sections of ***Result table***.

**Figure 11.**
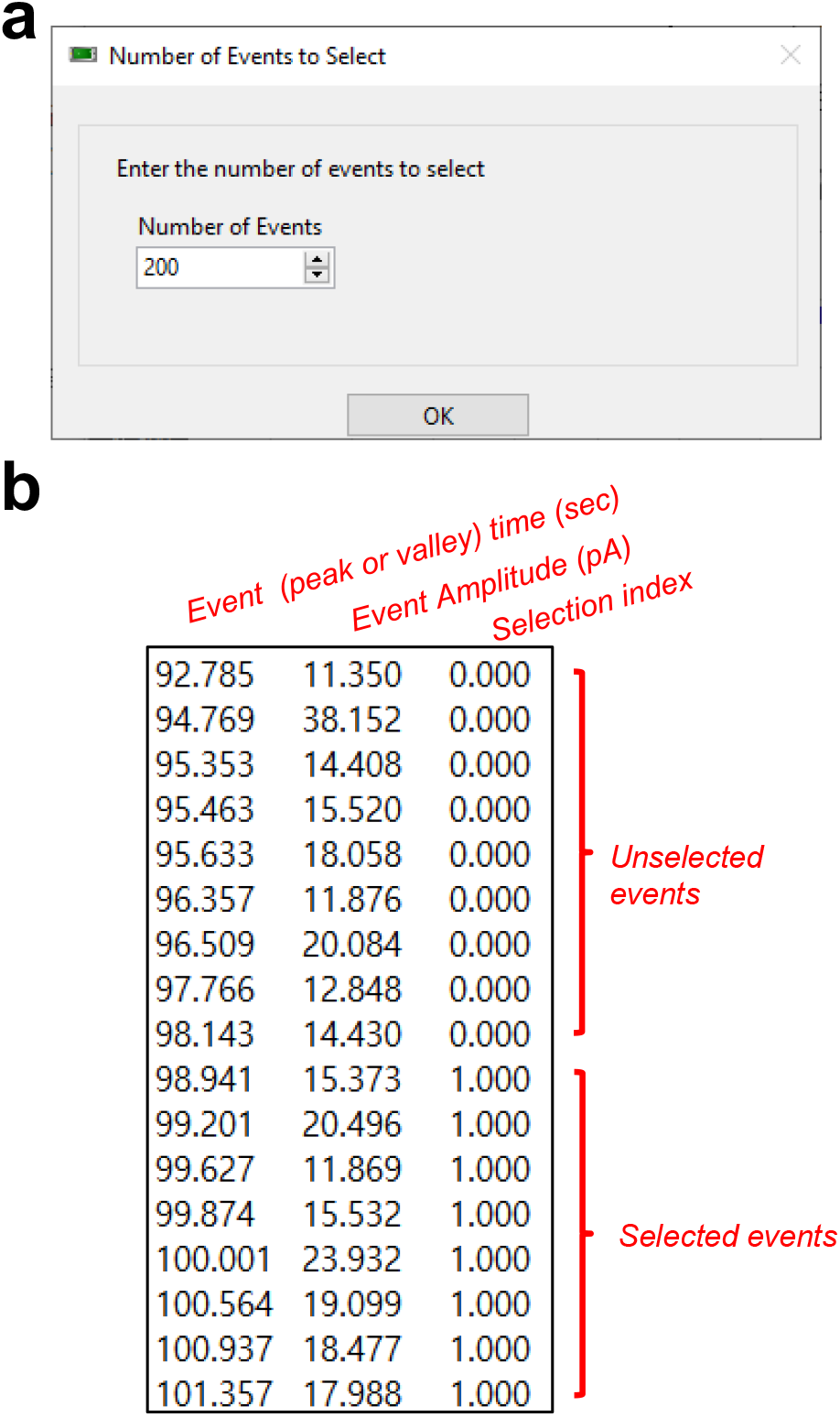
Selecting events and saving results in Minhee Analysis. (a) A pop-up window for entering the number of events to select. (b) A structure of unit file.

#### 4.2.6. Save result

To take advantage of the Minhee Retriever for the following steps, it is necessary to save the result of the event detection as a single file named unit file (*File>Save>Unit File (txt)*). The unit file is a tab-delimited text file which contains three rows without headers (Table 2 and Figure 11b). From the first row, it represents the event (peak or valley) time (sec), event amplitude (pA), and selection index. The selection index indicates whether an event has been selected in the selection process. Selected and unselected events are represented as the selection index 1 and 0, respectively. The selection index is used in the Minhee Retriever.

To maximize the benefits of file management using the Minhee Retriever, it is also recommended that a user uses the user’s own naming rule for the unit file. The Minhee Retriever searches unit files based on their names, which is described in detail in the use example of the Minhee Retriever below.

In addition, users can save the whole result (*File>Save>Whole Result (txt)*), including not only the results of the events but also the results of the baseline. Lastly, the Minhee Analysis automatically types its executing and analysis histories with a time stamp to ***Log*** window (Figure 6, (10)). A user can save the histories log as a text-formatted file (*File>Save>Analysis Log (txt)*).

#### 4.2.7. Miscellaneous features of Minhee Analysis

The Minhee Analysis has the following miscellaneous features:

***File>File explorer***. This function executes the Window File Explorer to show the currently analyzed file in the folder.
***File>Save>Raw trace (txt)***. This function allows users to save the raw trace of the currently displayed data in ***Sweep viewer*** as a text-formatted file.
***Edit>Copy to clipboard***. One simple way to export the result of the event detection is to copy the result to the clipboard and paste it into other third-party programs. This function allows users to copy the result to the Clipboard.
***View>Sweep Trace>Trace to display***. Users can choose between the raw and smoothed traces to be displayed in ***Sweep viewer***.

### 4.3. Use example of Minhee Retriever

#### 4.3.1. Retrieve the unit files

The Minhee Retriever has the function that a user can retrieve unit files in order to display merged results on the GUI of the Minhee Retriever. First, users specify the path of the folder a user will search for in ***Path of folder*** (Figure 7, (1)). Next, users may make a list of folder’s names in ***Folders to exclude*** (Figure 7, (4)) to specify folder names a user wants to exclude during the retrieval. Any folder a user excludes does not include in the retrieved results. This input is case insensitive. Then, users enter the pattern for files for which a user wants to search in ***Search pattern*** (Figure 7, (2)). Users can use the question mark sign (?) to match any single character. Lastly, a user executes ***Load*** function to search unit files that match with the filtering condition (Figure 7, (3))

To demonstrate the retrieval of unit files, we used the unit files which are provided by the package. Once the package is successfully installed in a PC, the example files can be found in the following path (C:\Program Files (x86)\NeuroPhysiology Lab\Example Files\Minhee Retriever). To generate these files, sEPSCs were recorded from the Purkinje cells (PC) of the Lobule III-V or Lobule X of cerebellar vermis (Figure 12a, left). All sEPSC data were analyzed by the Minhee Analysis. When we saved the files, our own customized naming rule was applied (Figure 12a, right).

**Figure 12.**
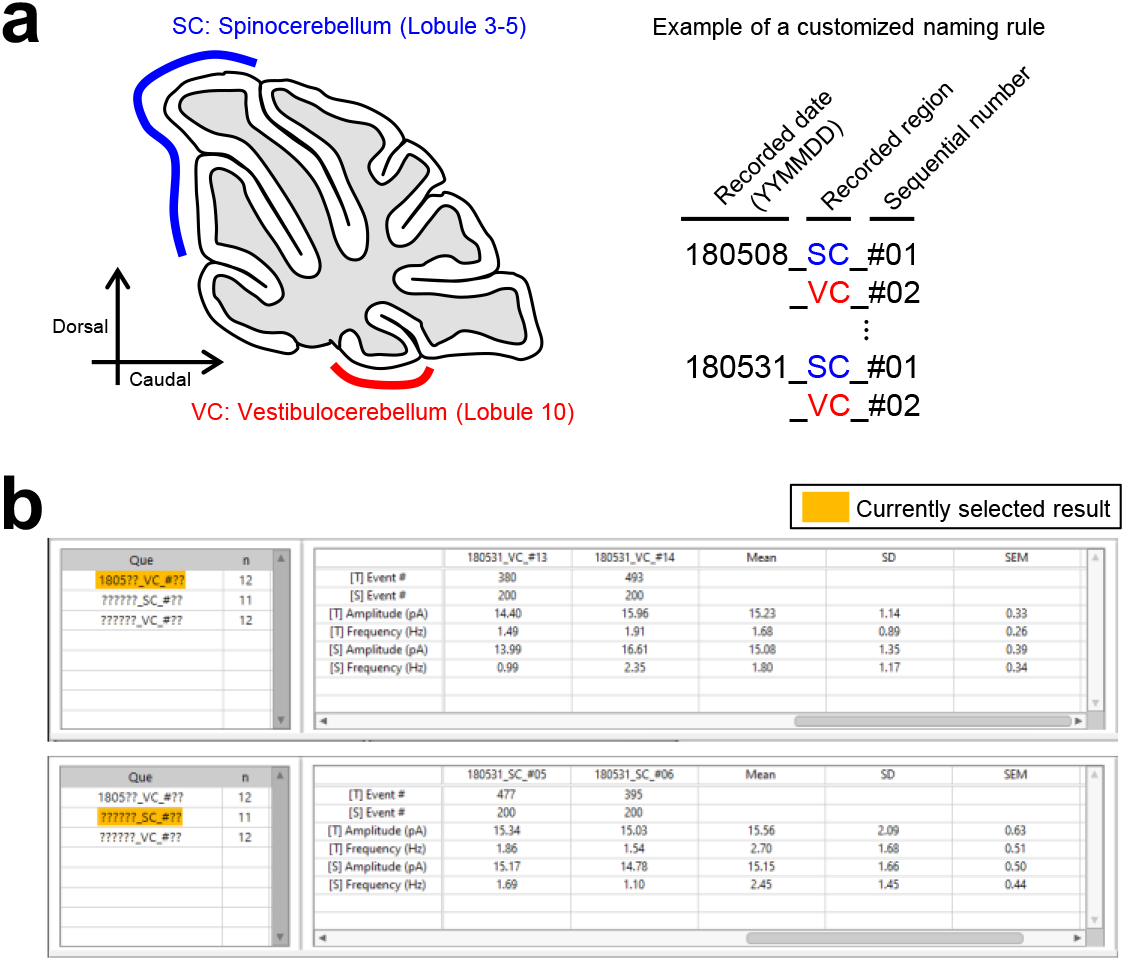
The search and browse functions for unit files in Minhee Retriever. Minhee Retriever allows users to retrieve and browse their unit files. (a) For example, if a user recorded sEPSC from the spino- and vestibulo-cerebellum (left), the file was named according to a customized naming rule (right). (b) After the retrieval of unit files is completed, a user can visualize the details by selecting an element from the stacked list.

The file name is composed of three codes, which is divided by the under-bar sign (_). In the first code, a six-digit code exhibiting a recorded date (YYMMDD) is placed. The code in the middle is a two-digit code standing for where the recording was made. For instance, SC and VC stand for recording from PCs of the Lobule III-V (spinocerebellum) and Lobule Ⅹ (vestibulocerebellum), respectively. The sequential number of cells written in the last code denotes the sequential order of the patch-clamp recording to assign a unique identification of the unit file and is placed after the sharp sign (#).

By naming like these, we took advantage of retrieving the unit files by a recorded date and/or the region where the recording was made (Figure 12b). For example, if we try to retrieve all files recorded from the vestibulocerebellum in May in 2018, we typed ‘1805??_VC_#??’ in ***Search pattern*** box. We also simply retrieved all files recorded from the spinocerebellum and vestibulocerebellum by typing ‘??????_SC_#??’ and ‘??????_VC_#??’ in ***Search pattern*** box.

#### 4.3.2. Browse retrieved data

Each time the ***Load*** function executes, the retrieved results will be stacked in the query table (Figure 7, (5)). Users can browse individual and averaged data from the retrieved files by selecting any item of the Que column in the query table. In the query table, the used search pattern and the number of retrieved files are shown in the first (Que) and second (n) columns, respectively. The result table (Figure 7, (6)) shows the number of events, amplitude, and frequency of the total and selected events in each unit file. In the header of the row, the t and s in the brackets ([t] and [s]) represent the total and selected events, respectively. Descriptive statistics such as mean, standard deviation (SD), and standard error of the mean (SEM) of the amplitude and frequency of individual files are calculated and added in the result table.

The Minhee Retriever also provides a generation of general histograms using the IEI and amplitude of selected events (Figure 7, (7)). Users can select elements of the retrieved list in the Que column to switch the data that a user wants to display. A user can initialize the whole retrieved list or delete the currently selected element by executing the ***Reset*** and ***Delete*** functions, respectively.

#### 4.3.3. Compare results between groups

The Minhee Retriever provides a comparative analysis between the retrieved results. First, users can compare the two distributions between experimental groups by generating a cumulative relative histogram of the IEI and amplitude of the selected events, which is the most widely used method for neurophysiologists to display PSC data [1, 2, 12, 14].

In this example, we retrieved unit files from four experimental groups: 180508_SC, 180508_VC, 180531_SC, and 180531_VC (Figure 13a). We simply generated two cumulative relative histograms of the IEI and amplitude of the events by executing the ***Cumulative plot*** function. Each histogram displays all retrieved groups currently listed on the query table. A user can specify the ***Number of bins***, and the ***Begin*** and ***End*** values for each histogram (Figure 7, (9)). A user can compare the relative distribution of two attributes between the groups.

**Figure 13.**
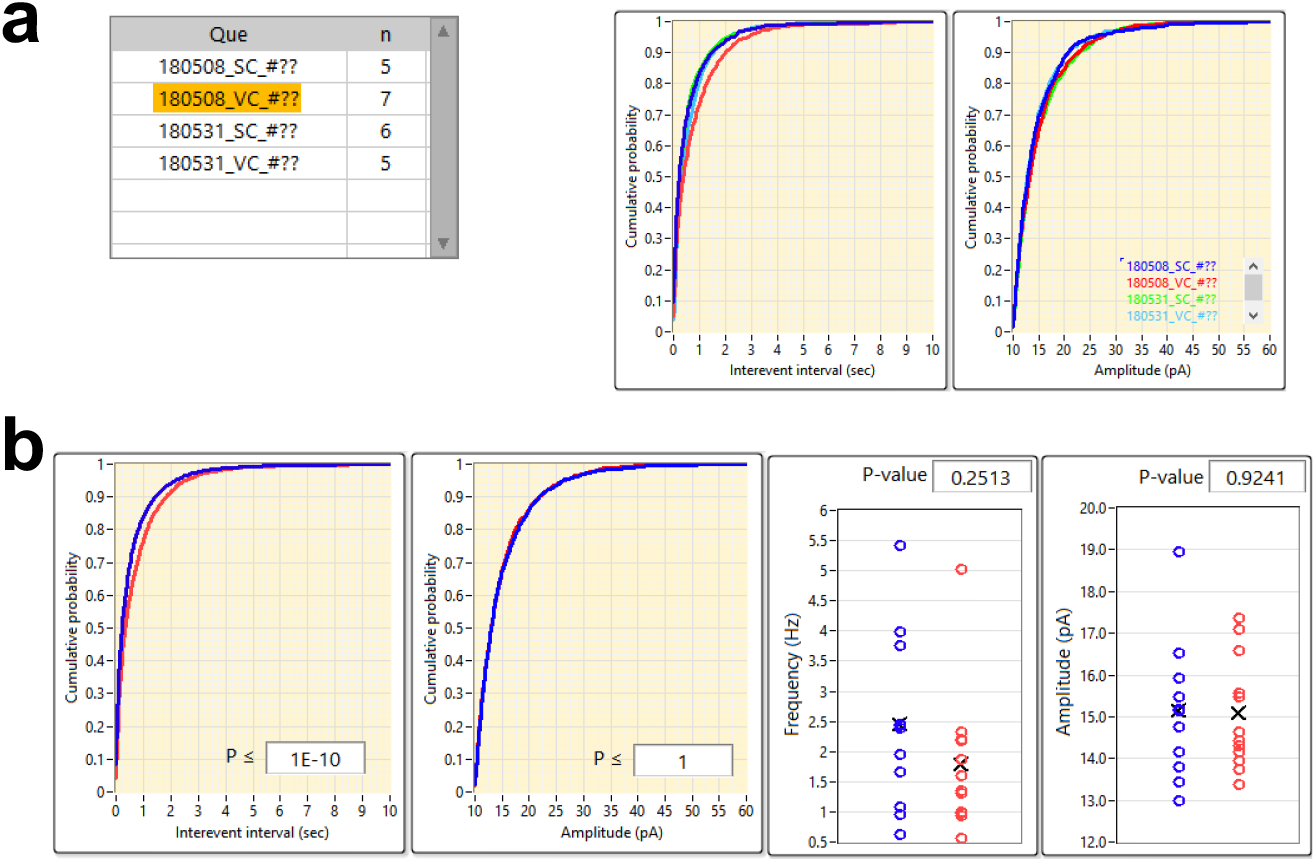
Comparison of results between the groups using Minhee Retriever. (a) Minhee Retriever provides a feature to generate cumulative relative histograms of the IEI and the amplitude of the events. Users can compare the relative distribution of two attributes among (right) all groups in the query table (left). (b) Users can test the statistical significance of the difference between the two groups. Minhee Retriever offers a Kolmogorov-Smirnov (K-S) test and independent two-sample t-test.

For testing the statistical significance of the difference between two groups, the Minhee Retriever serves two hypothesis tests, a kolmogorov-smirnov (K-S) test and an independent two-sample t-test. A K-S test is used to perform hypothesis tests in the relative cumulative distributions of the IEIs and the amplitude of events. An independent two-sample t-test is used to test whether the means of two attributes between two retrieved groups are equal or not.

We tested whether sEPSCs recorded in the spinocerebellum and vestibulocerebellum were drawn from the same statistical distribution. We retrieved all sEPSCs recorded in the spinocerebellum (n=11) and vestibulocerebellum (n=12) by entering ??????_VC_#?? and ??????_SC_#?? in the ***Search pattern***, respectively. Then, two data sets were set as the Data-set I and II, respectively (Figure 7, (10)). As a result of running the ***Comparison*** function, the p-value of the hypothesis test was obtained (Figure 13b). The cumulative distribution of the IEI of sEPSCs from the SC was shifted to the left, indicating that the sEPSC of the SC exhibited shorter intervals compared to those of the VC (p ≤ 1E-10, K-S test). However, in the comparison of the mean value, the difference in the IEI did not reach statistical significance (p = 0.2513, independent two-sample t-test). The cumulative distribution and mean values of sEPSC amplitudes were not different between the groups (p ≤ 1, K-S test; p = 0.9241, independent two-sample t-test).

#### 4.3.4. Miscellaneous features of Minhee Retriever

Minhee Retriever has the following miscellaneous features:

***Export***. This function generates a HTML file containing the cumulative relative frequency of the IEI and amplitudes.
***Save***. This function allows users to save the result table (Figure 7, (6)) as a text-formatted file.

## 5. DISCUSSION

Here, we presented an easy-to-use package that incorporates multiple steps of analysis of spontaneous synaptic events, from detection of PSCs to management of derived results in a row. Our algorithm for the detection of synaptic events was validated by using both simulated and experimental data. In addition, we described all the functionality of the package, general workflow for a successful analysis, and data management.

To decode the information encoded in the neural network, it is important to interpret the communication between neurons. Therefore, the measurements of PSCs have been utilized not only in several studies for the electrophysiological analysis of synaptic transmission in the nervous system [12, 14, 17], but also for the circuit mechanisms underlying certain types of behaviors [1, 13, 15]. Consequently, the importance of understanding the neuronal network has led to the development of novel tools to analyze PSC data [4–9].

The PSC detection method that we present in this study employs a stepwise exploratory algorithm based on the functions provided in the LabVIEW development environment, not an advanced function. We tested the validity of the algorithm by using both simulated and experimental data and compared results with those analyzed by other different PSC detection algorithms.

In addition to the event detection function, our package also provides a consecutive workflow that follows after event detection. Our method is to sort the analyzed results and allow neurophysiologists to make quick and efficient decision through visualization and statistical analysis. As mentioned above, PSC data analysis is considered as a primary measurement to understand the nature of neuronal conversations, therefore, our package is expected to be widely used to reduce the time and effort to analyze PSC data.

Another advantage of this package is that it provides an easy-to-use graphical user interface. Historically, many algorithms have proposed methods of detecting PSC [4–9], but from the end user’s point of view, whether the algorithm is well represented in an executable form is another important factor to consider [10, 18]. Since our package provides a graphical user interface for all features, it is easy for researchers (users) to use the package.

## Supplementary information

### Abbreviations

EPSC: Excitatory postsynaptic current
FN: False negative
FP: False positive
GUI: Graphical user interface
IEI: Interevent interval
IPSC: Inhibitory postsynaptic current
PSC: Postsynaptic current
TN: True negative
TP: True positive
VI: Virtual Instrument

## Acknowledgements

We thank Jewoo Seo for proofreading the manuscript.

## Funding

This research was supported by the National Research Foundation of Korea grants funded by the Korea government to SJK (NRF-2018R1A5A2025964, and NRF-2017M3C7A1029611).

## Availability of data materials

Minhee Analysis Package and data used in the present study are freely available to download from the repository: http://www.github.com/parkgilbong/Minhee_Analysis_Pack.

## Author contribution

YGK designed and developed the software. YGK and JJS performed experiments and data analysis. YGK, JJS, and SJK wrote and edited the manuscript. All authors read and approved the final manuscript.

## Ethics approval and consent for participation

Experimental procedures were approved by the Animal Care and Use Committee of Seoul National University.

## Consent for publication

Not applicable

## Competing interests

The authors declare that they have no competing interests.

## Footnotes

1 http://zone.ni.com/reference/en-XX/help/371361R-01/lvanls/sgfil/

2 https://zone.ni.com/reference/en-XX/help/371419D-01/lvasptconcepts/aspt_default_page/

3 http://zone.ni.com/reference/en-XX/help/371419D-01/lvwavelettk/wa_multiscale_peak_detection/

4 https://zone.ni.com/reference/en-XX/help/371361R-01/glang/recursive_file_list/

## Notes

https://github.com/parkgilbong/Minhee_Analysis_Pack

### Competing Interest Statement

The authors have declared no competing interest.

